# Quantifying the spatio-temporal image degradation under motion blur in fluorescence microscopy

**DOI:** 10.64898/2026.05.06.723301

**Authors:** Serafim Korovin, Kutay Ugurlu, Dylan Kalisvaart, Manon Kok, Rainer Heintzmann, Kirti Prakash, Carlas S. Smith

**Affiliations:** Delft Center for Systems and Control, Department of Mechanical Engineering, Delft University of Technology, Delft, 2628 CD, The Netherlands; Leibniz Institute of Photonic Technology, Jena, Germany; Institute of Physical Chemistry and Abbe Center of Photonics, Friedrich Schiller University Jena, Jena, Germany

## Abstract

The spatial resolution of optical imaging systems is fundamentally restricted by the diffraction limit. However, in widefield live-cell microscopy, the achievable resolution is further constrained by the specimen motion, which indicates the existence of a fundamental spatio-temporal resolution trade-off between signal accumulation during the full frame integration and the resulting motion blur. To improve the fidelity with which moving objects can be imaged, a quantitative understanding of this spatio-temporal trade-off is necessary. Here, we present a systematic analysis of motion-induced resolution dynamics measured with spectral signal-to-noise ratio (SSNR). We developed a simulation framework which models the image formation of objects undergoing arbitrary motion, to evaluate the degradation of the spatial resolution under translational and rotational dynamics. Our results demonstrate that for translating objects, the spatial resolution is anisotropically reduced as a function of the orientation of the object relative to the motion vector, leading to the spectral signal-to-noise ratio degrading by up to 50% and the resolution by up to 40% for a 90° change in the motion direction. Furthermore, we show that for rotational motion, conventional radially averaged metrics such as the Fourier Ring Correlation are not able to quantify the effects of angular blur. On the other hand, the SSNR is able to accurately quantify this degradation. These findings underscore the necessity of an object-oriented imaging approach, in which acquisition parameters such as exposure time are tuned to specific biological spatio-temporal characteristics to optimize the trade-off between motion blur and spatial fidelity.

## 1 Introduction

Over the last two decades, developments in microscopy have largely been driven by efforts to overcome the Abbe diffraction limit. Modern fluorescence microscopy achieves high molecular specificity, selectivity, and chemical contrast by using fluorophores that target specific biological samples [1]. With the introduction of super-resolution microscopy modalities, such as Structured Illumination Microscopy (SIM) [2–5], Stimulated Emission Depletion (STED) [6–9], and Single-Molecule Localization Microscopy (SMLM) [10–12], spatial resolution has successfully been extended down to the nanometer scale. Yet these spatial improvements fundamentally come at the cost of temporal trade-offs [13].Consequently, maximizing spatial resolving power fundamentally necessitates prolonged acquisition times [1, 14, 15].

This temporal cost introduces a severe bottleneck when imaging dynamic live-cell environments [1, 16, 17]. When the standard assumption of a static specimen is violated, the continuous-time emission signal is smeared across the detector during the extended integration time, irreversibly constraining the maximum recoverable spatial frequencies. This effect, known as motion blur, fundamentally degrades image fidelity. The resolution-constraining impact of motion blur becomes particularly problematic in fast-moving biological systems. For instance, highly dynamic cellular processes, such as organelle trafficking, frequently exhibit directional velocities reaching up to 50 nm ms^*−*1^. Assuming a standard 40 ms acquisition time and a 60 nm projected pixel size, the target structure undergoes a displacement of over 30 pixels within a single acquired frame. At the nanoscale, this substantial morphological distortion renders the resulting motion blur virtually indistinguishable from the object’s genuine biological structure.

To mitigate this spatio-temporal degradation, various hardware solutions have been developed. On the instrumentation front, extremely fast SIM systems have been developed that generate illumination patterns by switching between different fibers [18]. As a result, acquisition times can be improved to 112-140 frames per second. On the detection side, single-photon avalanche diode (SPAD) arrays have significantly advanced high-speed fluorescence lifetime imaging by enabling nanosecond-scale lifetime measurements [19].These advancements enable 100 FPS imaging in a parallel SPAD-array TCSPC fluorescence-lifetime imager [20], through highly parallelized detection with effectively zero readout noise and sub-nanosecond timing, thereby reducing required exposure times.

In addition to these hardware solutions, computational solutions have also been developed to reduce the spatio-temporal information loss. Particularly over the last few years, we have seen neural networks (NNs) transform low-resolution measurements into astonishingly super-resolved reconstructions of the cell [21–26]. Deep-learning frameworks such as CARE and DBlink have emerged to reconstruct super-resolved dynamics directly from low SNR data and motion-blurred single-molecule localization microscopy data, respectively, effectively bypassing the need for long temporal baselines [27, 28].

However, despite these advanced mitigation strategies, a quantitative understanding of how motion physically erodes the spatial frequency spectrum remains incomplete. While algorithms like DBlink effectively circumvent motion blur, thereby seemingly conjuring more information from less data, their black-box architectures obscure whether the recovered high-frequency details originate directly from the raw measurements or from learned priors embedded during network training [27]. To research the physical integrity with which these methods obtain a reconstruction, we need a quantitative and fundamental understanding of how motion blur affects spatial image frequencies. Additionally, to advance beyond empirical correction toward true object-oriented, adaptive imaging, it is critical to explicitly quantify the origin and degradation of spatiotemporal information [29].

Therefore, in this work, we systematically develop a quantitative framework to model how object kinematics fundamentally erode spatial resolution in fluorescence microscopy. By simulating benchmark targets undergoing both linear and rotational motion, we evaluate the anisotropic redistribution of spatial frequencies across varying exposure durations. Crucially, because conventional radially-averaged metrics like Fourier Ring Correlation are blind to the angular blur induced by rotation, we use a Spectral Signal-to-Noise Ratio (SSNR) metric dynamically calibrated against background variance. Through this approach, we aim to establish rigorous mathematical boundaries for motion blur, identifying the precise points at which photon accumulation shifts from constructive to destructive, thereby providing practical guidelines for preserving structural fidelity in dynamic imaging environments.

The rest of the paper is structured as follows: Section 2 describes the simulation procedure developed to facilitate our investigation, defines spectral signal-to-noise ratio (SSNR), and formalizes the SSNR-based frequency cut-off. The numerical simulation results are presented in Section 3, where we demonstrate the impact of representative motions on the resolution of various object types. We conclude with a discussion and an outline for future work in Section 4.

## 2 Methods

### 2.1 Image formation model

We built a computer program to study how motion changes image resolution. The full math description is in Supplementary Section 3. The program creates images in five steps.

First, we define the object’s shape and movement path. Second, we add a signal light to the object and a flat background light level to the whole space. Third, we put this model onto a fine grid of space and time. Fourth, we blur the object with a Gaussian function to model the microscope optics. Finally, we group the fine grid into camera pixels and exposure times. We add Poisson noise to this result to model the random detection of photons.

### 2.2 Image formation model

To quantitatively investigate how motion affects spatial resolution, we developed a simulator that models optical image formation for moving objects. We aimed to develop a time-dependent motion and shape discretization of an object with high fidelity to its continuous representation, and to simulate realistic camera measurements from them with a photon counting process represented by Poisson shot noise. The simulation pipeline consists of five main stages, which are briefly summarized in this subsection. An exhaustive description of the simulation process can be found in the Supplementary Section 3. The parameters used throughout our simulation are described in Supplementary Table 1.

We start by defining a continuous shape mask based on the intensity and background fluxes of the object. For this, we define a binary mask *S*(*x, y, t*) for continuous space-time coordinates (*x, y, t*), which takes the value 1 at space-time coordinates (*x, y, t*) if the object is present, and 0 otherwise. This definition includes both object geometry and motion-induced position changes. Then, the binary mask *S*(*x, y, t*) is scaled with a signal flux term *θ*_*I*_ (photons*/*(nm^2^ · ms)), representing the object intensity. To obtain the continuous object flux *o*(*x, y, t*) (photons*/*(nm^2^ ·ms)), a constant background flux *θ*_*bg*_ (photons*/*(nm^2^ ·ms)) is added to every space-time coordinate, which describes an expected intensity offset due to homogeneous background fluorescence. The model of *o*(*x, y, t*) is thus given by:

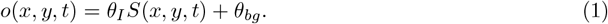

We then represent the continuous object flux on a finely discretized spatio-temporal mesh with voxel size *δV* = *δx δy δt* (nm^2^ · ms). The continuous object flux *o*(*x, y, t*) is sampled at coordinates corresponding to the mesh pixel centers (*x*^(*i*)^, *y*^(*j*)^, *t*^(*k*)^) for respective pixels at indices *i, j* and discrete time index *k*. Additionally, *o*(*x, y, t*) is multiplied with the mesh voxel size *δV* to preserve the total photon flux within the spatio-temporal voxel. As a result, we arrive at the following definition:

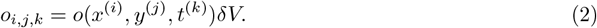

Next, we simulate a band-limited object representation on the image plane. To this end, the resulting discrete object *o*_*i,j,k*_ is convolved in space with the PSF of the simulated optical system. The PSF is represented as a Gaussian kernel with a standard deviation *σ* and a kernel support of *s* × *s* pixels. The Gaussian is a common choice for approximating the impulse response of optical systems, as it is both fast to compute and sufficiently accurate in 2D [30]. Additionally, our simulation shows that the implementation is numerically band-limited, with the spatial frequency at which the OTF amplitude drops below 10^*−*10^ × max(OTF) being 1/3 of the Nyquist frequency.

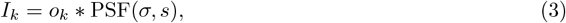

where ∗refers to the spatial convolution and *I*_*k*_ and *o*_*k*_ refer to the pixelated images at timestep *k*.

Finally, we simulate the camera readout process, which includes pixel discretization and photon counting during the exposure time, taken to be the inverse of the camera frame rate. We achieve that by binning the fine mesh pixels into camera pixels with factors (*C*_*x*_, *C*_*y*_) and binning each fine mesh frame with a factor *C*_*t*_. The resulting camera image *Î*_*p,q,r*_ for respective camera pixels at indices *p, q* and frame index *r* is defined as follows:

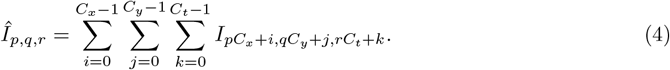

To model the photon-counting process, a shot-noise realization 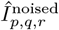 is sampled from a Poisson distribution with the expected value of *Î*_*p,q,r*_:

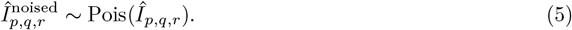

### 2.3 Estimation of spectral signal-to-noise ratio and resolution

We quantify image resolution using the spectral signal-to-noise ratio (SSNR). The SSNR is computed directly from groups of noisy image realizations to accurately reflect experimental noise characteristics without relying on assumed noise models. We express SSNR as ratio of mean spectra of the images to noise power computed as variance of the spectra of the realizations. It scales linearly with exposure time, which makes it more convenient to evaluate the change in resolution with varying exposure times. To ensure isotropic evaluation, the resulting SSNR values are averaged over all angles, yielding a direction-independent resolution metric. The detailed mathematical formulation for the batch-wise SSNR computation and radial averaging is provided in the Supplementary Section 4.

To determine the resolution limit from SSNR data, we must define a cutoff point. Standard thresholds from other methods, such as FRC, are not suitable to define a threshold because they do not align with SSNR value range. Instead, we define the resolution cut-off as the radial frequency at which the SSNR drops to the background SSNR level [31]. We also measure the total SSNR in a frequency band by integrating the SSNR curve. The mathematical procedure to obtain this threshold is provided in Supplementary Section 6.

## 3 Results

To quantify the effect of exposure interval on image formation and resolution, we conduct numerical simulations that isolate the impact of motion. We consider two basic geometries subjected to distinct motion types to illustrate these effects: a line undergoing translation and a Siemens star undergoing rotation. The line object offers a spectrum localized to one dimension, and the Siemens star, widely used for resolution assessment [32], provides rotational symmetry, which makes it particularly suitable for analyzing angular motion blur [33]. More on the object formation and motion types can be found in Supplementary Section 3.1.

This section is structured as follows: We first set a baseline by reporting the SNR in the spatial domain for a static Siemens star and SSNR for a static line. Next, we examine a rectangular object under translational motion and various orientations relative to the motion direction (Figure 3). We then analyze the Siemens star under rotational motion (Figure 4), where spectral modification induced by motion manifests mainly in the angular blur. The results are reported based on the relevant evaluations from Supplementary Table 2 for the quantification of Figure 2, Supplementary Tables 4 and 5 to provide numerical results for Figure 3, and Supplementary Tables 6 and 7 to quantify Figure 4.

### Static object imaging

In this section, we demonstrate and verify that the SNR increases with higher exposure time and intensity, and decreases with a higher background (as background is not part of the signal). Figure 1 presents a static Siemens star with eight spokes limited to a radius of 3.6 *µ*m imaged under varying exposure durations and background conditions. Figure 1*a* and *b* illustrate how the measurement is affected under varying background rates and exposure times, respectively.

**Figure 1.**
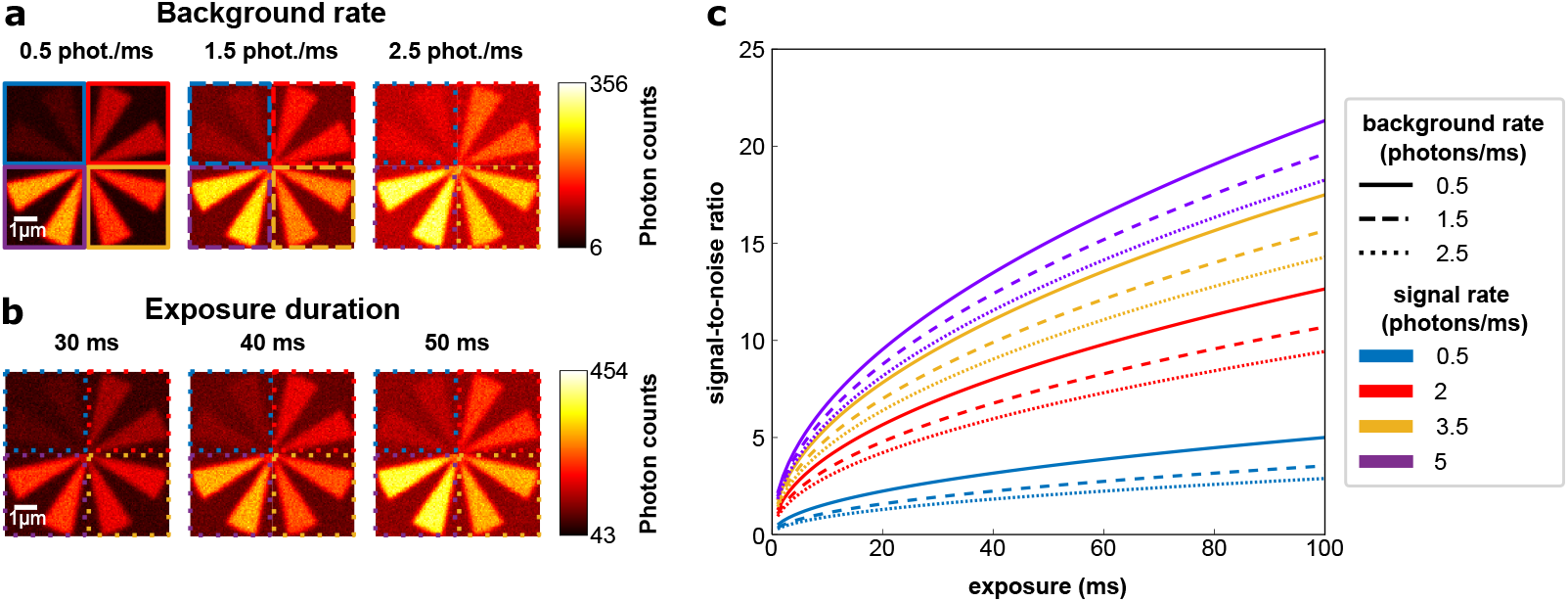
(*a, b*) A stationary Siemens star with varying *(a)* background rates imaged for 40 ms, and *(b)* exposure times at a background rate of 2.5 photons/ms. Each image is divided into color-coded quadrants, where each quadrant encodes the corresponding signal rate (in photons/ms), and the line type encodes the background rate (in photons/ms), as described in the legend. (*c*) Simulated signal-to-noise ratios across a range of exposure times for varying signal and background rates.

Figure 1*c* shows the maximum signal-to-noise ratio in each image computed using Equation (6) of Supplementary Section 1. Exposure time increases as the square root of the expected signal-to-noise ratio (SNR), while the SNR dependence on signal intensity is given by the factor 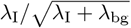. For example, raising the exposure time from 10 ms to 100 ms (with fixed signal and background) increases the SNR by a factor of 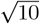. Moreover, increasing the signal rate from 0.5 to 5 for any fixed exposure time Δ*t*, increases this factor from 0.50 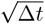 to 2.13 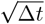, resulting in 4.26 times higher SNR (with a background rate of 0.50 photons). In contrast, increasing the background from 0.5 to 2.5 photon rate lowers the SNR from 0.5 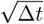 to 0.29 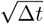 (a 42.40% decrease for a fixed signal rate of 0.5 photons). Notably, when the background photon rate is negligible (*λ*_bg_ ≪ *λ*_I_), SNR also exhibits a square root dependence on signal intensity. These numerical simulation results agree with the previous work [34, 35].

In the spectral domain, the SSNR displays a linear dependence on exposure time. This is displayed in Figure 2, which depicts a static line object with dimensions of 6 *µ*m in height and 0.12 *µ*m in width. Figure 2*a* presents a horizontal line imaged at increasing exposure durations, and the corresponding SSNR maps computed using Equation (39) are displayed in Figure 2*b*. A progressive increase in SSNR is observed with longer exposures, as reflected by the brighter intensities across the maps. The radially averaged SSNR, ⟨ SSNR⟩ (*ρ*), is shown together with the estimated background curve SSNR_bg_ and the cut-off frequency *ρ*_*c*_, which is identified by the cross in Figure 2*c*. For the exposure times of 10, 50 and 100 ms, we observe the cut-off frequencies increase from 3.86 to 4.31 and 4.47 *µm*^*−*1^ (averaged over angles using the values from Supplementary Table 2), respectively, which signifies the increase in the image resolution by 11.65% and 15.80% due to the signal accumulation. Moreover, the total SSNR measured by the area under the curve AUC(0, *ρ*_*c*_) rises from 1393.87 to 6956.51 and 13922.24 (averaged over angles) for the respective exposures, demonstrating a 399.08% (5-fold) and 898.82% (10-fold) increase in the total SSNR accumulated in the image.

**Figure 2.**
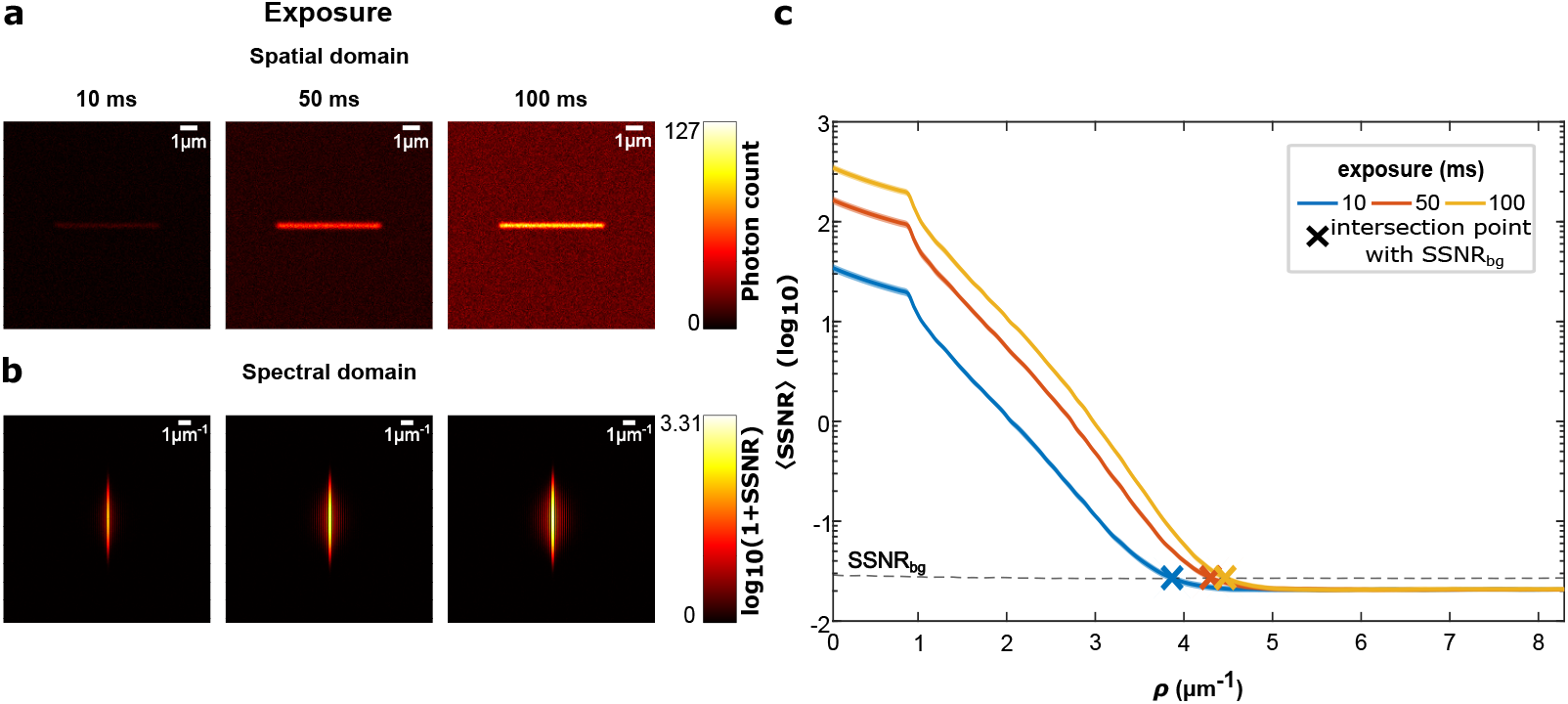
Increased exposure time improves SSNR for static objects. (*a*) Simulated images of a line object of height 6 *µ*m and width 0.12 *µ*m, oriented horizontally under the exposure times of 10, 50, and 100 ms. As the signal accumulates over time, the object appears brighter relative to the background. (b)The corresponding SSNR maps reveal that the spectral distribution aligns orthogonally to the line orientation. Due to linear dependence of SSNR on exposure time, the horizontal spectral component becomes increasingly dominant when the data is normalized and displayed on a logarithmic scale. (c)Radially averaged SSNR curves are shown for exposure times of 10 ms, 50 ms, and 100 ms. The frequency cut-offs, marked by cross symbols at the intersections of the SSNR curves with the background SSNR level (dashed black line), shift toward higher frequencies as exposure time increases. At the same time, the radial average curve increases across the whole bandwidth with the increase in exposure time.

To ensure consistency between computational and theoretical findings, we compare our numerical results to the derivations presented in Supplementary Section 2. When both velocity components *v*_*x*_ and *v*_*y*_ in Equation (15) are zero, there is no sinc modulation affecting the signal during static acquisition, and the SSNR increases linearly with exposure duration for all spatial frequency bands. Specifically, the peak magnitudes of SSNR at an exposure time of 100 ms display an increase of 0.30 and 1.00 on a logarithmic scale compared to 50 ms and 10 ms, respectively. These results correspond to the linear scaling factors of 2 and 10 for exposure times of 50 and 10 ms, respectively. Our numerical experiments verify this theoretical prediction, showing that, as exposure time increases, SSNR consistently rises and eventually surpasses the average background SSNR at higher frequencies, thus improving estimated resolution.

### Effect of translational motion on line object imaging

We simulated the line object of 6 *µ*m in height and 0.12 *µ*m in width from Figure 2 under various angular inclinations (0°, 45°, and 90° in the clockwise direction). For each line of different orientation, we applied translational motion across the *x*-axis to the object at a constant velocity *v*_*x*_ = 40 nm/ms, which matches the velocities observed in biological processes listed in Supplementary Section 3.6.

Figure 3*a* displays the accumulated images of the line for exposure times of 10 and 100 ms. As the exposure time increases, the line elongates horizontally along the direction of motion. For the 0° inclination, the line becomes thicker (6 *µ*m × 4 *µ*m), gradually approaching a square shape. For 45° inclination, the long exposure forms a parallelogram shape with a width of 4 *µ*m. For the 90° angle, the line width extended from 6 *µ*m to 10 *µ*m while the line height remains preserved. The resulting elongation occurs due to the accumulation of object snapshots during the translation along the motion direction. As a result, the motion of the 90° line results in a significant overlap of the signal photon accumulations over time, causing the visible extension to disappear. For the same reason, the 90° line reached the maximum photon count at 125 photons per pixel, compared to 46 and 47 photons per pixel for 0° and 45° lines, respectively.

**Figure 3.**
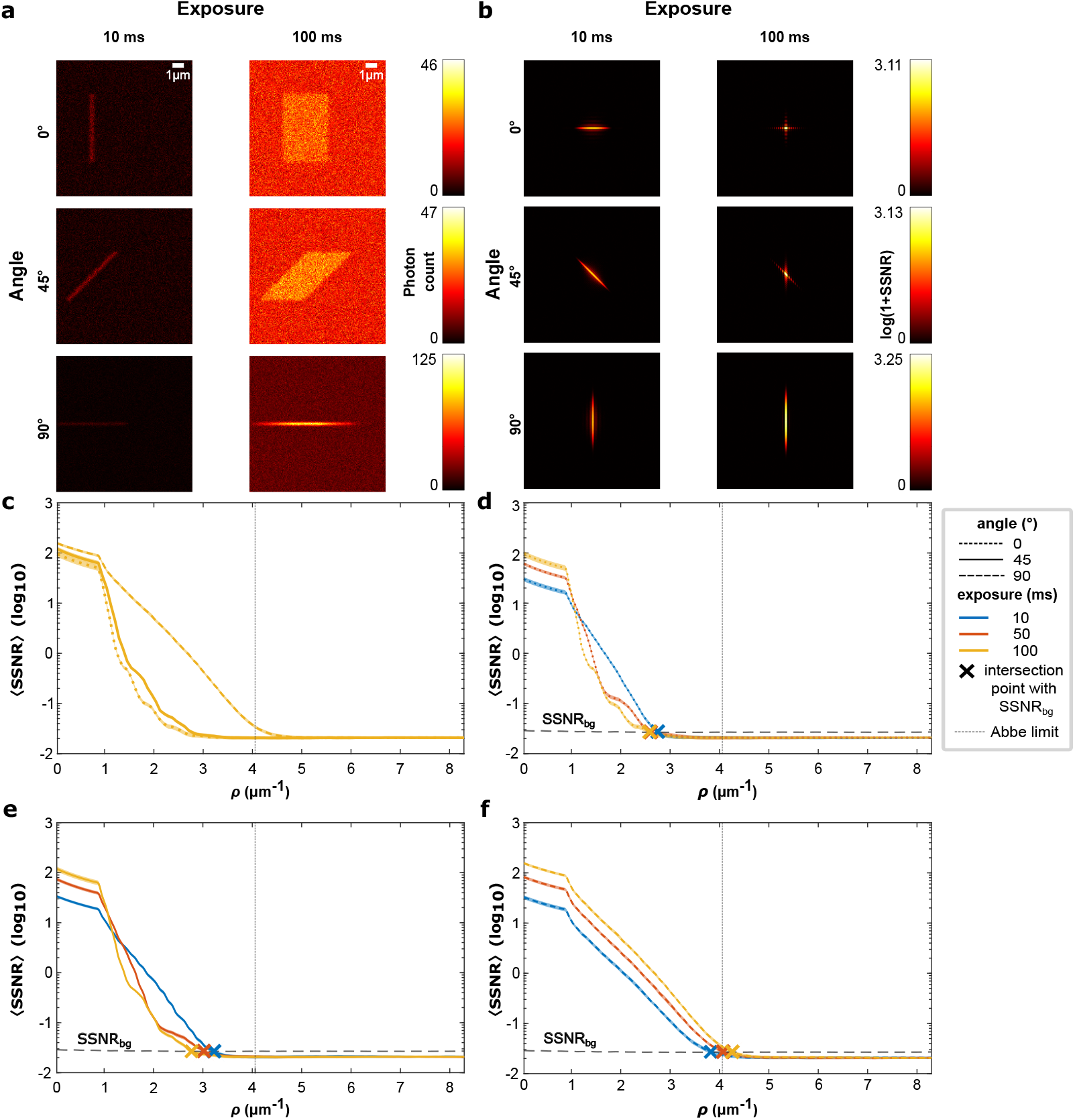
Translational motion blur on line objects with various inclinations. (*a*) Measured images of a line object oriented at 0°, 45°, and 90° under horizontal translation during acquisition. Two exposure times are shown: short (10 ms) and long (100 ms). The 0 and 45° lines exhibit a larger horizontal elongation after the long exposure time, while the 90° line preserves most of its structural integrity. (*b*) Corresponding SSNR maps indicate that horizontal motion broadens the spectrum along the vertical motion axis, most prominently for the 0° and 45° lines, whereas the spectrum for the 90° line remains relatively unchanged. (*c*) Radial averages of SSNR versus spatial radial frequency for long exposure time (100 ms). The 90° line demonstrates a higher radial average SSNR than that of the 0 and 45° lines. (*d-f*) Radial SSNR averages at varying exposures (10, 50, 100 ms) for 0° (in *d*), 45° (in *e*), and 90° (in *f*) angles. The 90° line, collinear to the motion, shows the greatest improvement in SSNR radial averages over increased exposures; the 0° and 45° lines show the redistribution of SSNR in the radial frequency spectrum with the increase in exposure time. Additionally, the gray vertical line depicts the Abbe diffraction limit which remains fixed with the change in exposure time.

In the frequency domain, the SSNR pixel maps (Figure 3*b*) illustrate the spectral modifications induced by motion. At an exposure time of 10 ms, the 0° line exhibits a spectrum concentrated along the *f*_*x*_ direction. When the exposure increases to 100 ms, a vertical frequency component emerges, indicating spatial blurring along the motion direction.

This effect stems from sinc modulation (see Supplementary Equation (15)), which suppresses frequencies aligned with motion and relatively emphasizes those in the orthogonal direction throughout the entire spectrum. A similar trend is observed for the 45° line: a single frequency component at 10 ms evolves into a dual-component structure at 100 ms. In contrast, the 90° line at 100 ms shows a constructive spectral effect: horizontal components diminish, and the intrinsic vertical spectrum of the object becomes more pronounced. This behavior results from the alignment between the object’s dominant spectral axis and the motion direction, as described by the sinc-shaped modulation function. Since only the *f*_*x*_ component is modulated (*v*_*y*_ = 0, see Equation (15) in Supplementary Section 2), the intrinsic spectrum of the line remains largely intact. The main spectral lobe, which contains 82.55% of the total SSNR and occupies 25% of the spectral domain vertically and 5% horizontally, is minimally influenced by this modulation.

As we examine the ⟨SSNR⟩ curves at a fixed exposure time of 100 ms shown in Figure 3*c*, we find that the 90° line contains the area under the curve (AUC) of 6190.45, compared to 3925.03 and 3065.40 for the 0° and 45° cases, respectively. This constitutes 36.60% and 50.48% loss of SSNR due to the respective orientation change. Similarly, the cut-off frequencies *ρ*_*c*_ = 2.58 *µ*m*−*1 for the 0° and 2.77 *µ*m*−*1 for the 45° line are 39.43% and 34.97% lower compared to the 90° line cut-off at 4.26 *µ*m*−*1, respectively. These results emphasize that the object orientation with regard to the translational motion direction can significantly degrade the signal-to-noise ratio by up to 50% and the image resolution by up to 40%.

Furthermore, as Figure 3*f* displays, the ⟨SSNR⟩ curve of the 90° line is *elevated* across the whole spectrum, with both *ρ*_*c*_ and AUC(0,*ρ*_*c*_) growing during the increase of the exposure times from 10 to 50 and 100 ms. Specifically, the frequency cut-offs rise from 3.84 to 4.08 and 4.26 *µ*m*−*1 (+10.94%), and the AUC(0,*ρ*_*c*_) increases from 1315.62 to 3273.22 and 6190.45 (+370.53%). Compared to the ⟨SSNR⟩ values obtained for a static line (Figure 2), the increase in frequency cut-offs under motion is lower than that of the static line (10.94% against 15.8%). Analogous results can be observed for the total SSNR, with an increase of 370.53% against 898.82%. These results highlight that motion blur degrades resolution even when the motion is predominantly aligned with the object orientation.

However, for 0° and 45° lines, increasing the exposure time *redistributes* the average radial frequency content contained in the ⟨SSNR⟩ (*ρ*), as visible in Figure 3*d* and *e*. Upon a visual inspection, we observe the presence of an “inflection frequency” *ρ*_infl_, wherein the ⟨SSNR⟩ (*ρ*) curves for various exposures intersect. We calculate the inflection frequencies as the minimum variance point between the ⟨SSNR⟩ (*ρ*) curves, and arrive to the *ρ*_infl_ values of 0.951 *µ*m*−*1 and 1.551 *µ*m*−*1 for 0° and 45° lines. We then compare the total SSNR for the frequencies below the inflection point with AUC(0, *ρ*_infl_) and above it with AUC(*ρ*_infl_, *ρ*_*c*_). With exposure times growing from 10 to 100 ms, we observe a consistent increase of 208.39% and 211.49% in AUC(0, *ρ*_infl_) for the respective 0° and 45° lines. In contrast, for frequencies above the inflection point, the AUC(*ρ*_infl_, *ρ*_*c*_) *decreases* by 28.48% and 78.59% for the same angles.

Moreover, we can observe from Figures 3*d*-*f* that the frequency cut-offs *ρ*_*c*_ increase with the expo-sure time for the 90° line, ranging between 3.84 *µ*m^*−*1^ and 4.26 *µ*m^*−*1^ (+10.94%) for the 10 and 100 ms exposures, respectively. For the 0° line, it falls from 2.76 *µ*m^*−*1^ to 2.58 *µ*m^*−*1^ (−6.52%), and for the 45° line, it decreases from 3.22 *µ*m^*−*1^ to 2.77 *µ*m^*−*1^ (−13.98%). This orientation-dependent pattern repeats the high-frequency total SSNR redistribution pattern, highlighting the resolution dependence on orientation, a property intrinsic to the object.

Additionally, we benchmarked SSNR-based resolution estimates with two-image and single-image FRC to verify the consistency of the results with other existing techniques. This comparison is presented in the Supplementary Tables 3 and 4.

### Effect of rotational motion on Siemens star object imaging

To investigate the influence of rotational motion on image resolution based on SSNR radial averages, we simulated a Siemens Star pattern with three spokes and applied clockwise rotational motion at angular velocities of 10.48, 6.98, and 5.24 radians/second, which correspond to rotation periods of 600, 900, and 1200 ms, respectively. These periods correspond to increasing angular displacement of 60, 40, and 30 degrees under 100 ms of exposure time. Figure 4*a* displays the accumulated images of the rotating Siemens star for exposure times of 10, 50, and 100 ms. With increased exposure time, the spokes become increasingly blurry, particularly at larger radial distances from the center of rotation, an effect commonly referred to as angular blurring or rotational motion blur [36]. This effect is further amplified by the higher angular velocities for a fixed exposure time.

**Figure 4.**
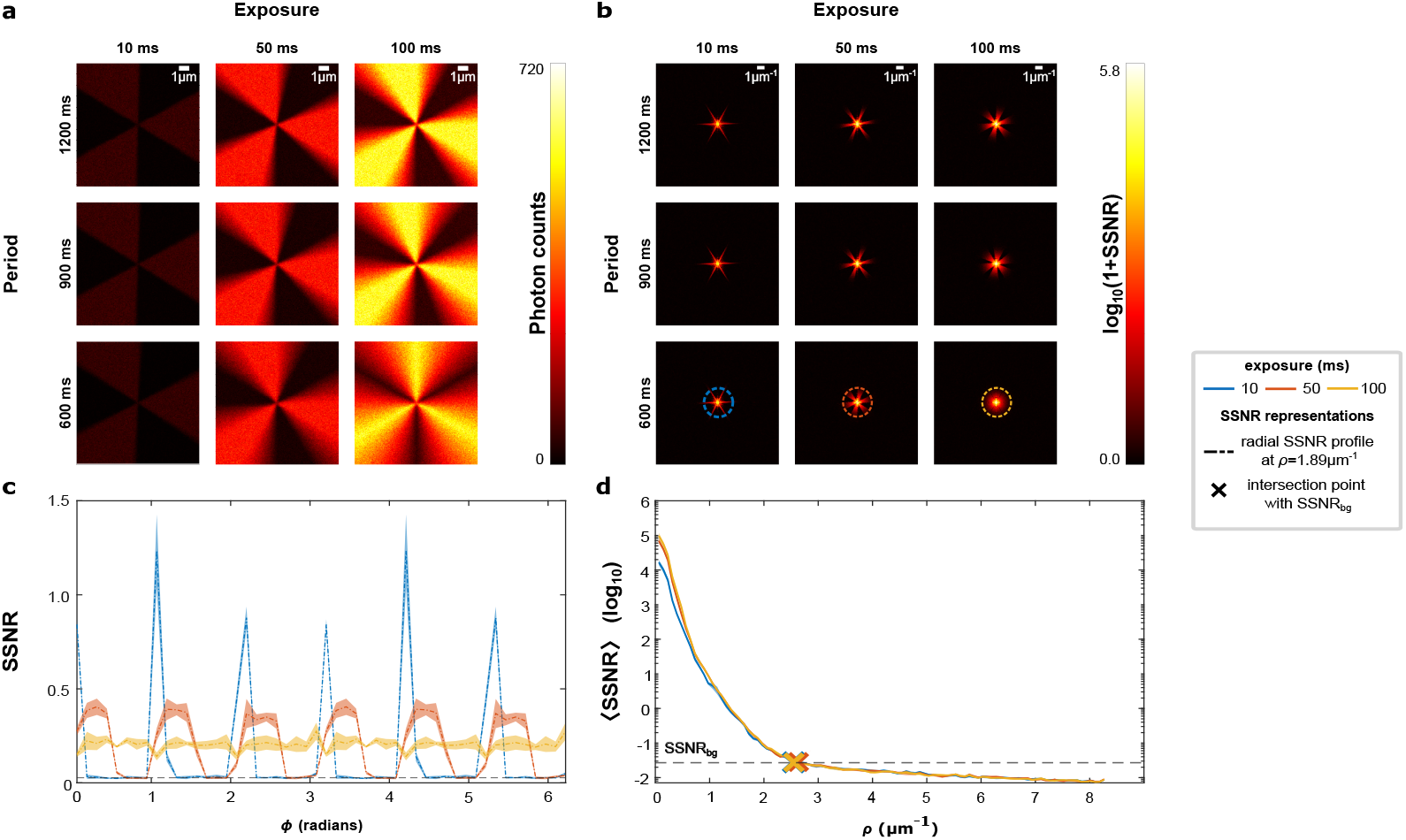
Rotational motion blur effects are concealed in SSNR radial averages, yet visible in radial profiles. (*a*) Reconstructed images of a Siemens star with three spokes rotating during acquisition with exposure times of 10 ms, 50 ms, and 100 ms. The angular blur of the spokes increases with both exposure duration and angular velocity (the inverse of the rotation period), due to the greater accumulated angular displacement during exposure. This effect scales with radial distance, producing more pronounced blurring at larger radii. (*b*) SSNR maps expose the angular dependence of this blur: SSNR in high-frequency regions, originally confined to narrow spokes, broadens in the angular direction with increasing exposure and angular velocity. (*c*) Radial slices (indicated by dashed circles in panel *b*) at *ρ* = 1.89 *µ*m^*−*1^ directly capture the angular blurring effect, with peaks broadening markedly as the exposure time increases. The dashed line in panel *c* shows the mean across multiple realizations, while the corresponding semi-transparent band indicates the interval spanning one standard deviation above and below the mean. (*d*) In contrast, full radial averages conceal the angular blur: all curves rise at low frequencies and converge to a similar cut-off, thereby masking the angular degradation evident in panels (a–c). For a stationary object, longer exposures would also increase the high-frequency SSNR; however, rotational motion suppresses this gain by redistributing SSNR across angles.

The frequency-domain representation displays a similar pattern. The SSNR pixel maps in Figure 4*b* display six narrow angular frequency components around the center for the exposure time of 10 ms. With an increase in exposure time and angular velocity, these components create motion blur in the clockwise direction. This demonstrates the spatial frequency redistribution effect caused by the rotational motion.

To quantify this effect, we investigated the SSNR radial profile at a fixed radial frequency *ρ* = 1.89 *µ*m*−*1, SSNR(*ρ* = 1.89*µ*m*−*1, *ϕ*), and a period of 600 ms (Figure 4*c*). At an exposure time of 10 ms, the profile rises to SSNR values of 1.25, followed by a rapid drop to a near-background SSNR value of 0.027 after an average non-background angular spread of 0.1676 radians, which constitutes 16.76% of the angular range. At 50 ms, the peak SSNR value decreases to around 0.35, but the angular spread broadens to about 0.5445 radians on average (52% of the angular range). For the 100 ms exposure time, the SSNR radial profile remains flattened across the entire 6.2832 radian range, centered at the SSNR value of 0.027, with no notable drops. Overall, we can observe an increase in the angular blurring with the exposure time.

In contrast, the radial average ⟨SSNR⟩ in Figure 4*d* shows an increase in AUC(0, *ρ*_*c*_) from 3.42 × 104 to 13.63 × 104 and 18.99 × 104 for the exposures of 10, 50, and 100 ms, accounting for 298.77% and 455.56% increases in total SSNR. The difference between the curves is predominantly pronounced for spatial frequencies below 1 *µ*m*−*1, resulting in a 455.76% increase in AUC for exposures from 10 to 100 ms. For the frequencies above 1 *µ*m*−*1, the total SSNR increases by just 12%, and no clear ordering of the cut-off frequencies can be observed as they yield values around *ρ*_*c*_ = 2.6 *µ*m*−*1 (±0.05 *µ*m*−*1). Therefore, it can be concluded that the rotational motion blur-induced SSNR redistribution is mostly concentrated in the lower frequencies, and thus cannot be detected by the cut-off frequencies alone.

Additionally, we explore how rotational velocity influences the resolution changes. For an exposure time of 100 ms, we observed the angular spread of the radial profile SSNR(*ρ* = 1.89 *µ*m*−*1, *ϕ*) increase from 0.59 for the period of 1200 ms to 0.71 for the period of 900 ms, constituting a growth of 0.12 radians. Furthermore, for an exposure time of 50 ms, the angular spread growth is smaller, at 0.08 (from 0.34 to 0.42 radian), for the same rotational periods. No measurable increase in angular blur was apparent at an exposure time of 10 ms (0.17 radians across all periods), suggesting that such brief exposure is sufficient to preserve the structural integrity of the object. Further details of this effect, shown across all exposure times and periods, are presented in Supplementary Table 7.

## 4 Conclusion

In this study, we have demonstrated that resolution is a dynamic variable governed by the fundamental trade-off between signal accumulation and object motion. By developing a simulation framework coupled with the SSNR metric, we quantified how these dynamics alter the information content of an image.

We systematically investigated how motion blur affects spatial information by simulating time-dependent object measurements and evaluating total SSNR and resolution cut-offs under dynamic conditions. First, we established a baseline by measuring SSNR for the static line object, where an exposure increase from 10 to 100 ms showed a ten-fold improvement in total SSNR and a resolution increase of 15.80%. However, under translational motion along the +*x* direction, lines oriented at 0° and 45° relative to the motion direction exhibit SSNR redistribution around an inflection frequency *ρ*_infl_: the SSNR is boosted by over 200% for the frequencies below *ρ*_infl_, and eroded by up to 80% for the frequencies above *ρ*_infl_.

Furthermore, we observe that object orientation influences the changes in resolution that occur with increasing exposure time. By increasing the exposure time from 10 to 100 ms, the cut-off frequency of the 90° line increases by 10.94%, but for the 0° and 45° lines, it falls by 6.52% and 13.98%. This effect reflects the spectral orientation dependence: 0° and 45° lines concentrate SSNR along the horizontal frequency direction *f*_*x*_ and thus suffer from a motion-induced sinc modulation, whereas the 90° line, whose spectrum is localized in the vertical frequency direction *f*_*y*_, remains largely unaffected. Notably, the increase in cut-off frequency observed for the 90° line represents an unusual scenario, in which motion has a constructive rather than degrading effect. We propose that this “constructive drift” phenomenon could be exploited as a novel, deliberate acquisition strategy in future research. For example, for highly anisotropic, parallel biological structures, such as actin filament bundles or striated muscle fibers, inducing a controlled stage drift parallel to the primary structural axis during acquisition could act as a physical low-pass filter, smearing isotropic background noise while actively accumulating constructive signal for the targeted structures. In typical imaging scenarios, however, object spectra are more isotropic rather than concentrated along a single axis. In such cases, motion attenuates high-frequency components, and the fine spatial details they encode are irretrievably lost due to motion blur. Due to the lack of structural priors, motion blur is indistinguishable from the object’s true shape, leading to misinterpretation of biological structures.

Additionally, our results highlight that the resolution metrics are strongly dependent on the intrinsic properties of the imaged object. We showed that varying a line orientation from 90° to 0° for a fixed translational velocity of 40 nm/ms in the +*x* direction at a fixed exposure of 100 ms resulted in the degradation of the total SSNR by up to 50% and of the image resolution by up to 40%. Furthermore, we demonstrated that increasing the rotational velocity from 1200 ms to 900 ms results in angular blurring increases of 0.08 and 0.12 radians at fixed exposures of 50 ms and 100 ms, respectively, but does not produce any significant changes at an exposure of 10 ms. As such properties cannot be directly controlled during real imaging scenarios, we recommend conducting an exploratory imaging of representative samples to obtain reliable estimates of the intrinsic parameters where possible. These estimates can be used to obtain a data-driven spatio-temporal prior, facilitating the detection of motion blur.

Crucially, our results expose a significant blind spot in standard resolution metrics. We found that increased exposure times cause severe angular blurring in rotating structures, where the SSNR radial profiles showed an increase in blurring from 16% of the angular range to 52% and 100% with the exposure increases from 10 to 50 and 100 ms. However, we also demonstrate that conventional radially averaged metrics, such as FRC, fail to detect this degradation, since they mask the angular motion blur. This creates a risk of having false confidence, where a researcher may believe they have achieved high resolution based on the metric, while the true underlying biological structure (e.g., a rotating bacterial flagellum) is severely degraded. Therefore, more work is required to develop resolution metrics for arbitrary motions, and we strongly recommend utilizing radial SSNR profiling rather than radial averages when analyzing rotationally dynamic specimens.

Minimizing movement-induced image degradations requires a methodological shift from solely maximizing SNR to optimizing the spatio-temporal trade-off between exposure duration and the motion blur in the sample. To achieve this, we suggest transitioning toward object-oriented, adaptive imaging paradigms. Recent advancements, such as the hybrid Event-Driven Acquisition (hybrid-EDA) framework [29], have demonstrated the viability of using low-phototoxicity phase contrast surveillance to monitor cellular dynamics in real-time. By synthesizing our SSNR framework with such AI-driven multimodal hardware, future microscopes could continuously estimate the velocity vectors of a target structure using label-free surveillance. Upon detecting rapid movement, the system’s onboard AI would calculate the exact inflection frequency *ρ*_infl_ and instantly trigger high-intensity fluorescence lasers with an exposure duration strictly tuned to remain below this destructive spatio-temporal threshold.

This object-specific acquisition protocol balances SNR against motion blur, ensuring that photon accumulation remains entirely constructive. By establishing these absolute mathematical boundaries for kinematic degradation, our work provides a critical physical safeguard for structural fidelity, ensuring that super-resolved biological dynamics represent genuine optical measurements rather than hallucinated artifacts from computational over-reconstruction.

In practical terms, we recommend adopting an object-specific imaging approach based on our results to obtain accurate resolution estimates under dynamic movements. Object-specific properties, such as velocity estimates, can be obtained in an exploratory imaging step utilizing label-free phase contrast surveillance to avoid unnecessary photobleaching. While doing so, we highlight the need to adjust resolution metrics to incorporate angular information rather than averaging it, thereby enabling more accurate resolution evaluation under rotational motion. We recommend employing a two-step acquisition protocol: an initial high-speed, low-SNR exploratory scan to estimate velocity vectors, followed by an acquisition where exposure time is tuned to the specific inflection frequency of the sample. Using this a priori knowledge with the adjusted resolution metrics will help determine an acquisition speed that balances SNR against motion blur. As a computational alternative to balancing SNR and exposure time, a promising direction could be to simultaneously image in separate slow, high-SNR and fast, low-SNR regimes, with subsequent computational extraction of relevant information. Such empirical exploration, utilizing dual-exposure priors, can guide the identification of optimal operating points that are specific to the imaging system without the need for an initial exploratory imaging step, and could push the spatial and temporal boundaries of imaging fast biological processes.

## Additional information

### Data and code availability

The code that supports the findings of this study is openly available on GitHub [37] at https://github.com/qnano/spatio-temporal-analysis.

## Author contributions

S.K., K.U., D.K., K.P., and C.S.S. designed the research. S.K. and K.U. derived the metric, implemented the experiments, analyzed the data and wrote the initial draft manuscript. D.K. derived the simulation model. R.H. contributed discussion and ideas. The draft manuscript was edited by all authors. The study was supervised by C.S.S.

## Acknowledgments

S.K., K.U., D.K., and C.S.S. were supported by the Netherlands Organisation for Scientific Research (NWO), under NWO Vidi project no. 20390.

## Declaration of interests

The authors have no conflicts to disclose.

## Supplementary Citations

The following citations appear in the supplementary file: [32, 38–58].

## Notes

### Competing Interest Statement

The authors have declared no competing interest.

